# Non-redundant roles for the human mRNA decapping cofactor paralogs DCP1a and DCP1b

**DOI:** 10.1101/2023.10.19.563039

**Authors:** Ivana Vukovic, Samantha Baranda, Jonathan W. Ruffin, Jon Karlin, Steven B. McMahon

**Affiliations:** Thomas Jefferson University, Department of Biochemistry and Molecular Biology, Philadelphia, PA, 19107, USA

**Author notes:** Corresponding author: Steven B. McMahon.

## Abstract

Eukaryotic gene expression is regulated at both the transcriptional and post-transcriptional levels, with disruption of regulation contributing significantly to human diseases. In particular, the 5’ m7G mRNA cap is the central node in post-transcriptional regulation, participating in both mRNA stabilization and translation efficiency. DCP1a and DCP1b are paralogous cofactor proteins of the major mRNA cap hydrolase DCP2. As lower eukaryotes have a single DCP1 cofactor, the functional advantages gained by this evolutionary divergence remain unclear. Here we report the first functional dissection of DCP1a and DCP1b, demonstrating that DCP1a and DCP1b are distinct, non-redundant cofactors of the decapping enzyme DCP2, with unique roles in decapping complex integrity and specificity. Specifically, DCP1a is essential for decapping complex assembly and for interactions between the decapping complex and mRNA cap binding proteins. In contrast, DCP1b is essential for decapping complex interactions with the translational machinery. Additionally, DCP1a and DCP1b impact the turnover of distinct mRNAs. The observation that different ontological groups of mRNA molecules are regulated by DCP1a and DCP1b, along with their non-redundant roles in decapping complex integrity, provides the first evidence that these highly similar paralogs have qualitatively distinct functions. Furthermore, these observations implicate both DCP1a and DCP1b in transcript buffering.

## Introduction

mRNAs represent a distinct functional RNA group whose regulation is a critical aspect of eukaryotic gene expression. Most mature mRNAs in eukaryotic cells are capped by 7-methylguanosine at their 5’ terminus (m^7^G) (1–3). The mRNA cap controls critical stages of the mRNA lifecycle: recruiting complexes involved in mRNA processing, mRNA export, translation initiation, protecting mRNA from degradation by 5’-3’ exoribonucleases, and marking mRNA as “self” to avoid being targeted by the innate immune system (1,4). Since the mRNA cap has these essential roles in the mRNA lifecycle, precise regulation of mRNA capping and decapping is critical (2,5,6).

mRNA capping occurs co-transcriptionally through three distinct enzymatic reactions (6–8). RNA triphosphatase first remove the 5’ y-phosphate from the nascent pre-mRNA to generate a 5’ diphosphate end (7–9). RNA guanylyl transferases then cap the pre-mRNA with GMP (7–9). Finally, RNA guanine-N7 methyltransferases transfer a methyl group from the S-adenosylmethionine methyl donor to the N7 position of the terminal guanine base (7,10,11). Cap binding complex (CBC) binds to the mRNA cap and recruits proteins which facilitate further mRNA processing, i.e. splicing and polyadenylation, before mRNA is exported into the cytoplasm (1,12,13). The cap binding proteins protect mRNA from degradation by physically blocking the cap from being bound and hydrolyzed by the decapping enzymes (14). In the cytoplasm, mature mRNA can have several different fates: 1) they can be translated, 2) they can be held in processing bodies (PBs) in a translationally repressed state, or 3) they can be degraded (15). The binding of the eIF4F complex, consisting of the initiation factors eIF4E, eIF4G, and helicase DDX6, to the mRNA cap initiates mRNA translation (12,16,17). PBs are cytoplasmic non-membranous ribonucleoprotein granules that contain high concentrations of RNA and RNA binding proteins (18–20). Decapping complex members and exoribonucleases are among the proteins localized to PBs, leading to a model in which PBs are also a site of mRNA decapping and subsequent degradation (18,20,21). More recently, it has been discovered that PBs store translationally repressed mRNAs (21,22). This translational repression of mRNA associated with decapping can be reversed, providing another point at which gene expression can be regulated (21).

As with all other aspects of mRNA metabolism, degradation of mRNA is tightly regulated (23,24). While the bulk of cellular mRNA is degraded through decapping, followed by 5’-3’ degradation mediated by the exoribonuclease XRN1, some mRNAs are degraded by the exosome and a 3’-5’ degradation pathway (4,23,25). Degradation is initiated by the shortening of the polyA tail by deadenylases (15,26). Human cells contain a family of NUDIX proteins, several of which have intrinsic mRNA decapping activity in vitro. Among the NUDIX proteins which hydrolyze the m^7^G mRNA cap are DCP2/Nudt20 and Nudt16 (4,27) The N-terminus of human DCP2 contains a regulatory domain (RD), followed by a catalytic domain (CD) (28), while the C-terminus is intrinsically disordered (29). The CD consists of a conserved NUDIX hydrolase domain which hydrolyzes diphosphates linked to nucleosides (28,29). The CD and RD domains of DCP2 are linked by a flexible hinge and undergo a rapid transition between an open and closed state (28–30). Due to its flexible nature, defining the structure of the active and inactive conformations of DCP2 has been a challenge (28,29,31). Most structures of DCP2 utilize the Schizosaccharomyces pombe or Kluyveromyces lactis proteins (29). DCP2 has low basal decapping activity in vitro, and its enzymatic activity is greatly enhanced by its interaction with its main decapping activator DCP1 (29,32). DCP1 contains an Ena/Vasp Homology domain 1 (EVH1 domain), known to mediate interactions of multiprotein complexes and to transduce migratory and morphological signals into the cytoskeleton, at its N-terminus. DCP1 proteins also contain a Trimerization Domain (TD) at their C-terminus (33). DCP1 binds to DCP2 through an asparagine–arginine containing *loop* (NR loop) in the EVH1 domain, an interaction that converts DCP2 conformation from an inactive/open to an active/closed state (28). The human genome encodes two DCP1 paralogs, termed DCP1a and DCP1b. DCP1a and DCP1b can both homo- and hetero-trimerize via their TD domain (TD), and it is the trimeric form of DCP1a/DCP1b that is found with DCP2 in the decapping complex (34). In human cells, the interaction between DCP2 and DCP1 is weak and is stabilized by the binding of the decapping enhancer protein EDC4 (28). From a functional perspective, efficient in vivo decapping activity requires the binding of EDC4 to DCP1 and DCP2, and EDC4 depletion leads to an inhibition of decapping (28).

To date, mechanistic studies have focused primarily on the single DCP1 protein in yeast and the DCP1a paralog in humans. Currently we lack an understanding of the overlapping and unique aspects of DCP1a and DCP1b function. The studies reported here broaden our understanding of mRNA decapping in general by demonstrating that DCP1a and DCP1b indeed have unique roles in human cells. We report that while DCP1a is integral in decapping complex integrity, both DCP1a and DCP1b foster unique interactions between decapping complex subunits. Specifically, DCP1a is responsible for maintaining interaction with RNA cap binding proteins and DCP1b for interactions with the translational machinery. DCP1a and DCP1b also impact the stability of distinct sets of mRNAs. For example, DCP1a controls mRNAs that encode proteins with a role in adaptive immunity and transcription, while a separate group of DCP1b-dependent mRNAs also plays a role in transcription. These findings provide the first functional dissection of the distinct roles played by the human DCP1 paralogs in mRNA metabolism.

## MATERIALS AND METHODS

### Cell culture, lentivirus production, cell infection, transfection and CRISPR

Human cells (HCT116 and HEK293T) were obtained from American Type Culture Collection (ATCC) and cultured in Dulbecco’s modified Eagle’s medium (DMEM, Corning) supplemented with 10% fetal calf serum (FBS, Gemini Bio-Products) at 37 °C in 5% CO2. Cells were tested at four week intervals for mycoplasma (Sigma-Aldrich). All stock plates were treated with plasmocin (InvivoGen) as per manufacturer’s instructions.

Packaging plasmids (psPAX2 + pMD2.G) were co-transfected into HEK293T cells, along with shRNA or cDNA expression vectors, to deplete or overexpress DCP1b or DCP1a. Viral supernatants were collected, filtered, and added to the appropriate cells in the presence of 8 µg/ml polybrene (Sigma Aldrich). Puromycin (Sigma Aldrich) was used for selection of infected cells.. For knockdown, HCT116 cells were infected with lentiviral carrying shRNA specific to DCP1a (OriGene) or DCP1b (OriGene) and for overexpression with lentiviral expression plasmid for DCP1b (OriGene). As a control luciferase shRNA was used, obtained from TRC collection (Sigma-Aldrich).

To delete the DCP1b or DCP1a genes, HCT116 cells were transfected with the appropriate mixture of 3 different sgRNAs (Synthego Inc.) and Cas9 protein (Synthego Inc.), as per manufacturer’s instructions. Briefly, Cas9 was diluted to 3 µM with Opti-Mem (ThermoFisher Scientific) and sgRNAs were diluted to 3 µM with TE buffer (Synthego Inc). The RNP complex was assembled by incubating Cas9 protein, sgRNAs and at a ratio of 1.3:1 for 10 min at room temperature. Transfection solution was prepared by incubating 3.5 µl Lipofectamine CRISPRMAX (Thermo Fisher Scientific) and 25 µl Opti-Mem per sample for 5 min at room temperature. RNP was added to the transfection solution and the mixture incubated for 20 min at room temperature. Cells were trypsinized, washed with PBS, and counted. 10^5^ cells were used per transfection reaction. As a control, sgRNA was omitted from the RNP complex assembly. Cells were cultured for a total of 72 hours at which time editing efficiency of the pooled cells was determined by PCR amplification of the appropriate genomic region, followed by DNA sequencing of the PCR product. Polyclonal pools were identified at this point. Single cell clones were isolated, cultured and validated, using genomic PCR, DNA sequencing and western blotting.

### Western blotting and Co-immunoprecipitation Assays (Co-IP)

In preparation for western blotting and co-immunoprecipitation assays, cells were washed twice with PBS and harvested. Cell lysis was performed for 15 min on ice in an E1A buffer (20 mM NaH2PO4, 150 mM NaCl, 50 mM NaF, 0.5% (w/v) IGEPAL, 2.5 mM EDTA, 125 mM sodium pyruvate, and 10% (w/v) glycerol) with proteinase inhibitor cocktail (1:1000, Milipore Sigma). Lysates were centrifuged at 20,000 x g for 10 min at 4 °C and protein concentrations were determined using Pierce BCA Protein Assay Kit (ThermoFisher Scientific) according to manufacturer’s instructions. For western blots, inputs for Co-IPs, sodium dodecyl sulfate (SDS) loading buffer (250 mM Tris–HCl (pH 6.8), 8% (w/v) sodium dodecyl sulfate, 0.2% (w/v) bromophenol blue, 40% (v/v) glycerol, 20% (v/v) β-mercaptoethanol) was added to the lysates which were boiled for 7 min. For Co-IPs and IPs that were analyzed by LC/MS/MS, lysates were incubated with the appropriate antibody, and equivalent amount of the control IgG antibody, at 4 °C overnight. Total amount of protein (per one IP) for DCP2 was 300 µg, DCP1a/1b 270 µg and for DDX6 300 µg. Protein A/G beads (Santa Cruz) were added to the lysates for 1 hr at 4 °C to capture the precipitate. The lysates were centrifuged for 5 min at 3,500 RPM. The beads were washed 3x with E1A buffer supplemented with proteinase inhibitor cocktail. SDS loading buffer was added to lysates which were boiled for 7 min.

Lysates were separated by the SDS-polyacrylamide gel electrophoresis (SDS-PAGE) and electroblotted onto nitrocellulose membranes (BIO-RAD). Membranes were blocked for 30 min in 5% dry milk (AppliChem) reconstituted in TBS-T [20 mM Tris, 150 mM NaCl, 0.1% Tween (Millipore Sigma)] and incubated with the appropriate antibodies overnight at 4 °C (DCP1a, DCP1b, DCP2, DDX6, EDC4, EDC3 1:1000 in TBS-T, GAPDH 1:500 in TBS-T). The membranes were washed 3 x 10 min in TBST and incubated with the secondary anti-mouse or anti-rabbit antibodies for 1 hr at room temperature. ECL Western blotting solution (FisherScientific) was used to visualize bands and KwikQuant imager (Kindle Biosciences) was used to develop the images.

### LC/MS/MS analysis

SDS gels were stained with Colloidal Blue (Invitrogen). These stained regions were excised and reduced with TCEP, iodoacetamide was used for alkylation, and finally, digested with trypsin. Tryptic digests for DDX6 IP, DCP1a IP and DCP1b IP were analyzed using the standard 90 min LC gradient on the Thermo Q Exactive Plus mass spectrometer. Tryptic digests for control, DCP1a KO and DCP1b KO proteome cell analysis, were analyzed using an extended 4h LC gradient on the Thermo Q Exactive Plus mass spectrometer. MS data were searched against the Swiss Prot human proteome database (DDX6 IP: 6/4/2021, DCP1a IP, DCP1b IP and proteomics study of control, DCP1a KO and DCP1b KO cells: 10/10/2019) using MaxQuant (DDX6 IP: 1.6.17.0, DCP1a and DCP1b IP:1.6.3.3., proteomics study for control, DCP1a KO and DCP1b KO cells: 1.6.15.0) Protein and peptide false discovery rate was set to 1% and fold changes were calculated using intensity values.

For DDX6 IP, three replicas were performed and subjected to the analysis pipeline described above. Proteins were considered high confident if they were 1) minimum absolute fold change of 2, 2) q-value < 0.05, 3) identified by a minimum of 2 razor + unique peptides in one of the sample groups compared, and 4) detected in at least 2 of the replicates in one of the sample groups compared. Normalized intensity (as per DDX6 intensity) was used to generate heatmaps for DDX6 IP (Figure 2B, 2C).

**Figure 1.**
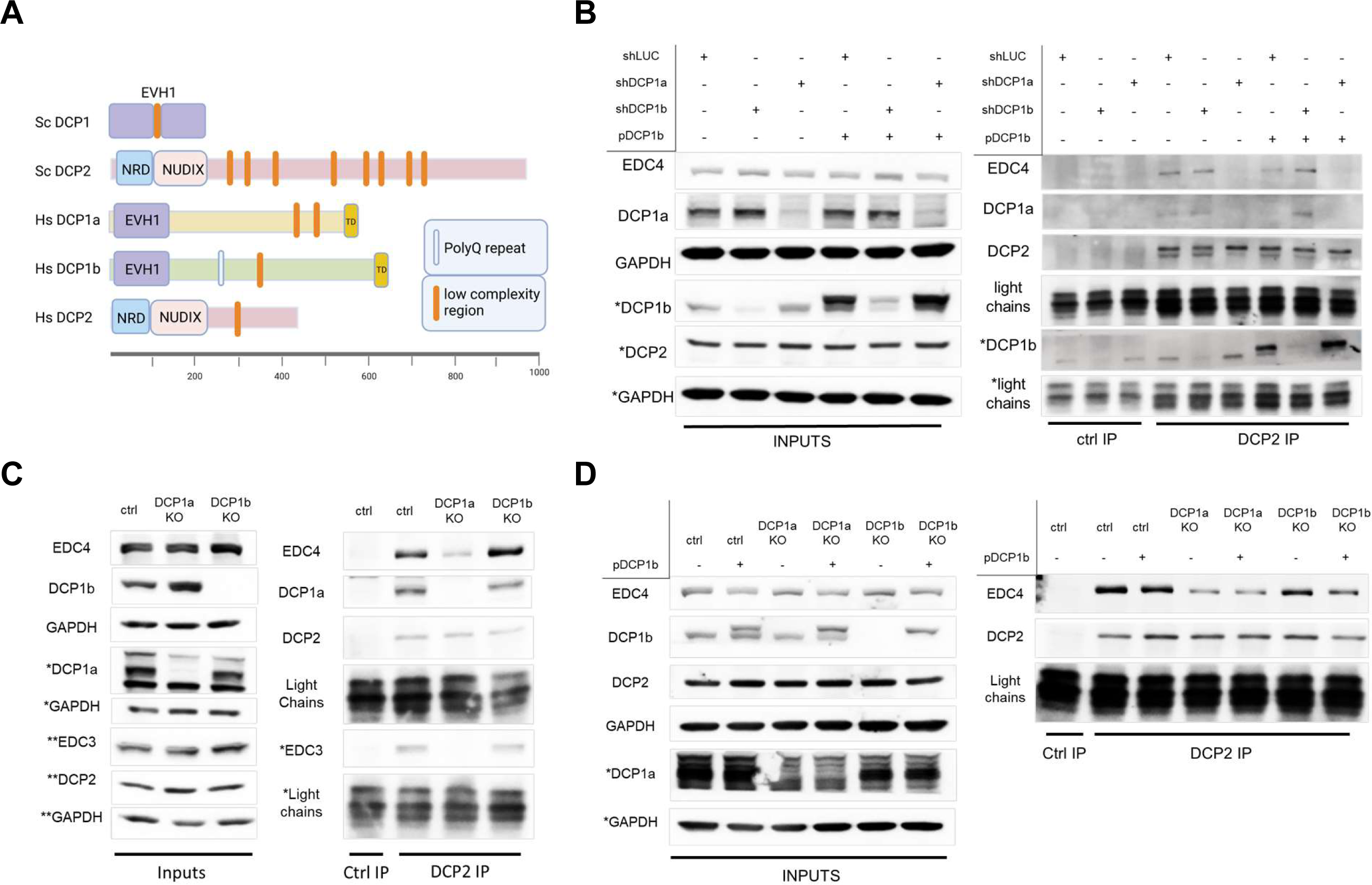
DCP1a, and not DCP1b, mediates the interaction of DCP2 with EDC4 and EDC3. **A)** Comparison of DCP2 and DCP1(a/b) protein domains in *Saccharomyces cerevisiae* (Sc), and in *Homo sapiens* (Hs). EVH1 - Ena/Vasp Homology *domain* 1, NRD -N-terminus by an a-helical regulatory domain, TD – trimerization domain. *The protein domains were defined using* http://smart.embl-heidelberg.de/*. Created with BioRender.com. **B)*** HCT116 wt cells were infected with lentiviruses carrying shRNAs targeting LUC, DCP1b or DCP1a. On day two, the cells were selected with puromycin. On day five, cells were infected with the second virus expressing DCP1b or an empty vector control. Cells were harvested three days following the second round of infections and DCP2 was immunoprecipitated. Western blots (WB) showing inputs, IgG and DCP2 immunoprecipitation in shLUC, shDCP1a and shDCP1b cells. **C)** HCT116 cells were transfected with Cas9 protein and no gRNA to create control cells or Cas9 protein and a mix of sgRNAs targeting either DCP1a or DCP1b. Three days post transfection single cell clones were selected and cultured for ten days. Clones were validated and DCP2 was immunoprecipitated. WB showing inputs, IgG immunoprecipitation in control, DCP2 IP in Control (ctrl), DCP1a KO and DCP1b KO cells. **D)** Ctrl, DCP1a KO (polyclonal pool) and DCP1b KO (single cell clone) cells were infected with a lentivirus expressing DCP1b or the empty vector control. Three days post-infection cells were harvested and DCP2 was immunoprecipitated. WB show inputs, IgG and DCP2 IP.

**Figure 2.**
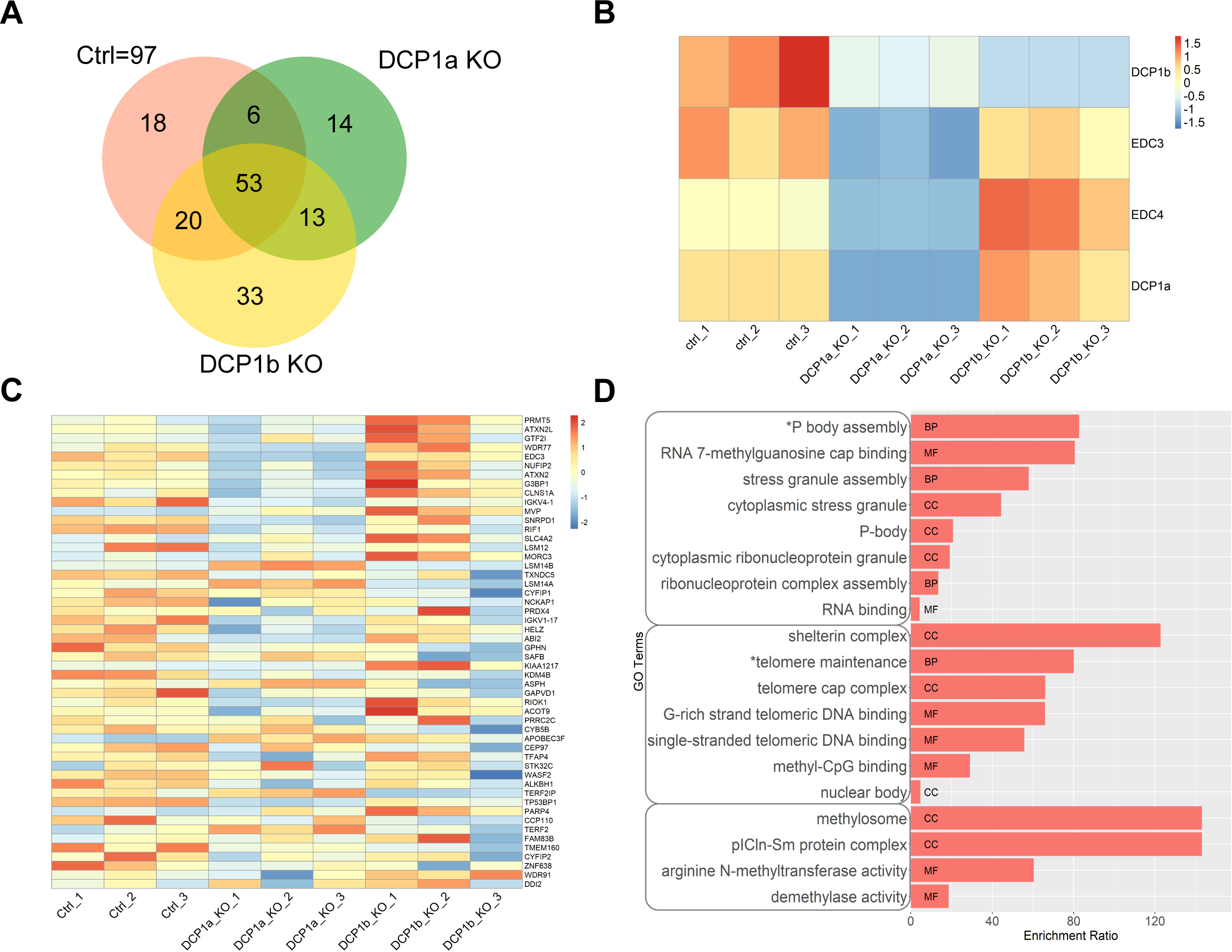
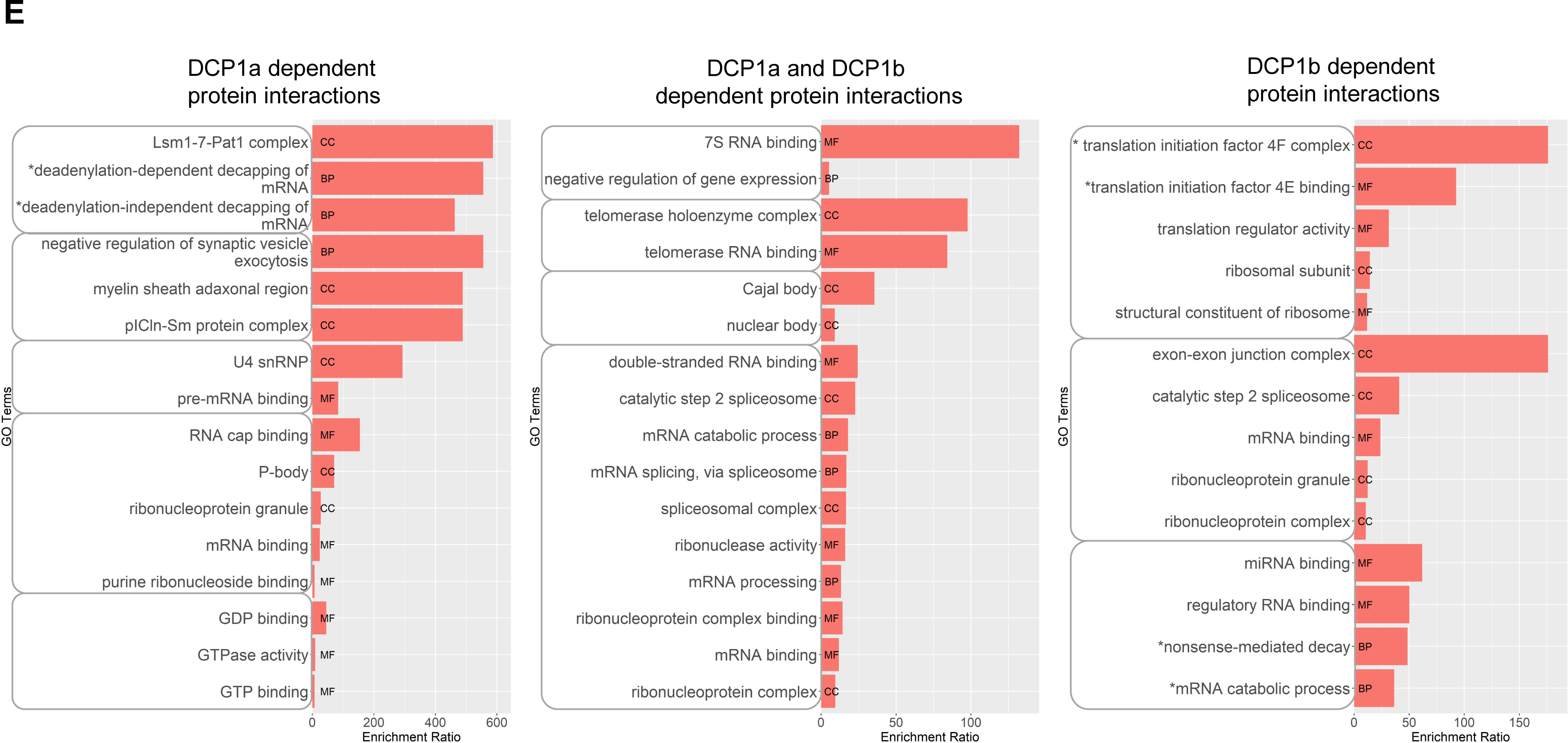
DCP1a is critical for interactions between known decapping complex members. **A)** DDX6 was immunoprecipitated in control cells, DCP1a KO and DCP1b KO cells with IgG immunoprecipitation as a control. IPs were sent for profiling by LC/MS/MS. Venn diagram represents the high confident proteins found in all conditions. **B)** Heatmap reports intensities (sum of peptide peaks as analyzed by mass spectrometry) of the core decapping complex proteins DCP1a, DCP1b and EDC3 in the DDX6 IP, for each condition. Biological replicas 1, 2 and 3 are shown. The heatmap is scaled by row. **C)** Heatmap showing intensities of the core 53 proteins, which do not depend on DCP1a or DCP1b presence or absence, in DDX6 IP. Biological replicas 1, 2 and 3 are shown. The heatmap is scaled by row. **D)** Ontology analysis of the core 53 proteins present in DDX6 IP. GO terms with similar cellular role are grouped. Only relevant GO terms with p value < 0.05 are shown. MF-Molecular function, BP – Biological process, CC – Cellular Compartment. **E)** Ontology analysis of the DDX6-interacting proteins dependent on DCP1a (6 proteins), DCP1b (20 proteins), and proteins dependent on both DCP1a and DCP1b presence (18 proteins). GO terms with similar cellular role are grouped. Only relevant GO terms with p value < 0.05 are shown. MF-Molecular function, BP – Biological process, CC – Cellular Compartment.

For DCP1a and DCP1b IP, two replicas were performed and proteins were considered high confidence if they were 1) identified by a minimum of 2 razor + unique peptides in the experimental condition, 2) identified in both experimental replicas and 3) have minimum fold change of 2 (if not identified in the IgG control) or 5(if the protein is also identified in the IgG control).

Proteins were considered high confidence in ctrl, DCP1a KO and DCP1b KO cells if 1) p value <0.05, 2) identified by a minimum of 2 razor+ unique peptides in any of the samples compared and 3) detected in at least 2 of the triplicates in any of the groups compared.

Please see supplementary files with DDX6 IP, DCP1a IP, DCP1b IP and proteomics study of ctrl, DCP1a KO and DCP1b KO datasets.

### SLAM-seq/GRAND-SLAM

SLAM-seq Kinetic Kit was purchased from LEXOGEN and the manufacturer’s instructions were followed. Briefly, cells were cultured to appropriate density and media containing 100 µM 4SU was added. The cells were cultured for 0 hr and 4 hrs, while maintained in a light-free environment. Cells were lysed at times indicated using TRIZOL and lysates subsequently frozen at −80 °C. All RNA isolation steps were performed in light-free environment. RNA concentration was determined using NanoDrop (Implen) and 5 µg of RNA was alkylated by incubation at 50 °C for 15 min, with a mixture of 100 mM iodoacetamide, 25 μl of Organic Solvent and 5 μl of Sodium Phosphate. After the reaction was terminated, light-free conditions were no longer employed. Ethanol precipitation was performed to isolate the RNA according to manufacturer instructions (LEXOGEN). mRNA libraries were prepared using the QuantSeq 3’ mRNA-Seq Library Prep Kit FWD (Illumina). Quality of mRNA libraries was determined using Agilent Tape Station and mRNA was sequenced at 75 bp single read sequencing using NextSeq 500 (Illumina).

### Dot Blot Analysis of 4SU labeled RNAV

4SU incorporation was confirmed with dot blot analysis of 4SU labeled RNA, prior to alkylation. 4SU labeled RNA was biotinylated with EZ-link Iodoacetyl-LC-biotin (Pierce), resulting in irreversible biotinylation as described (35). Briefly, 350 µl (50 µg of RNA diluted in nuclease-free H2O) of 4SU labeled RNA was incubated with 50 µl 10x Biotinylation Buffer (100 mM Tris pH 7.4, 10 mM EDTA in nuclease-free H2O) and 100 µl of EZ-link Iodoacetyl-LC-biotin (1 mg/ml in dimethylformamide) at room temperature for 1.5 hr while rotating in dark conditions. RNA was isolated by chloroform extraction and in 2 ml Phase Lock Gel Heavy tubes to reduce the loss of RNA. RNA was precipitated with 5 M NaCl and isopropanol to the water phase, centrifuged and the precipitate was washed with ethanol. Zeta membrane (Biorad) was incubated in nuclease-free H2O while rocking for 10 min and air-dried for 5 min. 5 µl of 200 ng/µl RNA dilution in ice cold dot blot binding buffer (10 mM NaOH, 1 mM EDTA) was applied to the Zeta membrane by pipetting. Biotinlyated, non 4SU RNA was used as a control. The membrane was air-dried for 5 min, incubated for 30 min in 40 ml blocking buffer (20 ml 20% SDS with 20 ml 1x PBS pH 7-8, and EDTA to the final concentration of 1 mM) with rocking, and then incubated with 10 ml of 1:1000 streptavidin-horseradish peroxidase for 15 min. Membrane was washed twice in 40 ml PBS + 10% SDS (20 ml PBS + 20 ml 20% SDS) for 5 min, twice in 40 ml PBS + 1% SDS (38 ml PBS + 2 ml 20% SDS) for 5 min, and twice in 40 ml PBS + 0.1% SDS (40 ml PBS + 200 μl 20% SDS) for 5 min. Residual buffer was removed by blotting and membrane-bound HRP was visualized as described above.

### Data Analysis for Slam-seq dataset

SLAM-DUNK, in conjunction with the globally refined analysis of newly transcribed RNA and decay rates using SLAM-seq (GRAND-SLAM), was used to analyze SLAM-seq datasets (36,37). SLAM-DUNK is an automated SLAM-seq data analysis pipeline and features MultiQC plugin for diagnostic and quality assurance purposes (36,37). SLAM-DUNK output was used to run GRAND-SLAM. SLAM-DUNK v0.4.3 was used and the following code was used to run this pipeline:

> slamdunk all -r /mnt/e/slamseq_run_9_9_2021/hg19_no_alt_analysis_set.fa -b /mnt/e/slamseq_run_9_9_2021/pure_UTR_3_hg19_ensemble.bed -o /mnt/e/slamseq_run_9_9_2021/ /mnt/e/slamseq_run_9_9_2021/sample_file.tsv

GRAND-SLAM output includes the proportion and the corresponding posterior distribution of new and old RNA for each gene. GRAND SLAM v2.0.6 version was used and the following code was used to run the pipeline:

> gedi -e Slam -genomic homo_sapiens.75 -prefix test2/24h -progress -plot -D -full -reads bams.bamlist SLAM-DUNK can be found at https://t-neumann.github.io/slamdunk/ and GRAND-SLAM at https://github.com/erhard-lab/gedi/wiki/GRAND-SLAM

### Half-life estimation

GRAND-SLAM output “MAP” was used as the new-to-total RNA ratio (NTR), and background was subtracted (0 hr timepoint serving as the background). Half-lives were estimated using the formula Half-life = -T* ln2/ln(1-NTR), where T is the time that the cells were labeled with 4SU. The dataset has three replicas, Z score was used to remove outliers and at least two replicas were used to estimate the final half-lives. The maximum half-life estimation was set to 24hrs, and variance of at most 0.3 was allowed to consider each half-life estimate significant. The max half-life estimation was set to 24hrs, as a 4hr labeling window can not accurately estimate very long half-lives. Please see supplementary file for dataset with half-lives estimates.

### Cell Viability assays

Cells were cultured to an appropriate confluence and media containing 0 µM, 50 µM, 100 µM or 200 µM 4SU was added. Cells were cultured in the presence of 4SU for 6 hrs while supplying fresh 4SU media every 3 hrs. At 6hrs, media was replaced with DMEM without 4SU and 24hrs later cell viability was evaluated. Culture media was collected, and cells were harvested for evaluation of cell death. Cells were washed with PBS and stained with either nothing or with propidium iodide (PI) and/or Annexin V. Cell death was assessed with CytoFLEX S Flow Cytometer (Beckman Coulter).

### Ontology analysis

WebGestalt (WEB-based Gene SeT AnaLysis Toolkit) is a functional enrichment analysis web tool that was used to examine ontology of different sets of proteins or mRNAs (38). The overrepresentation analysis method was used, reference set was genome, top 10 categories for Molecular Function, Biological Process and Cellular Component were combined and analyzed. Only the relevant categories with p value < 0.05 were represented in the bar charts, full lists are provided as supplemental data.

### Cellular fractionation

Cellular fractionation assay was performed as previously described (39).

### Reagents

#### Antibodies

DCP1a (Abcam, Cambridge, UK, ab47811), DCP1b (Cell Signaling Technologies, Danvers, Massachusetts, USA, mAb #13233), EDC4 (Abcam, Cambridge, UK, ab72408), EDC3 (Cell Signaling Technologies, Danvers, Massachusetts, USA, mAb #14495), GAPDH (Cell Signaling Technologies, Danvers, Massachusetts, USA, mAb #5174), DDX6 (Novus Biologicals, Littleton, Colorado, NB200-192)

#### Enzymes

SpCas9 2NLS Nuclease (Synthego Inc, Redwood City, CA, no cat number)

#### Kits

Gene Knockout Kit v2 - human - DCP1A - 1.5 nmol (Synthego Inc, Redwood City, CA, no cat number)

Gene Knockout Kit v2 - human - DCP1B - 1.5 nmol (Synthego Inc, Redwood City, CA, no cat number)

SLAMseq Kinetics Kit – Anabolic Kinetics Module (Lexogen, Vienna, Austria, 061.24) SLAMseq Kinetics Kit – Catabolic Kinetics Module (Lexogen, Vienna, Austria, 062.24) QuantSeq 3’ mRNA-Seq Library Prep Kit FWD for Illumina – (Lexogen, Vienna, Austria, 015.24)

Mycoplasma PCR Detection Kit (LookOut, Sigma-Aldrich) Colloidal Blue Staining Kit (Invitrogen, Carlsbad, CA, LC6025)

#### Non-standard chemicals

Plasmocin (InvivoGen, ant-mpp) Lipofectamine plus (Invitrogen, CMAX0003) Zeta membrane (Biorad, 162-0153)

### Biological Resources

HEK293T, Human Cell line, ATCC: CRL-3216™

HCT116, Human Cell line, ATCC: CCL-247™

HCT116 derived cas9 control cell line (M17, M4)

HCT116 derived DCP1a KO cell line (single cell clone A3, polygenic clone A9)

HCT116 derived DCP1b KO cell line (single cell clone B2, single cell clone B18)

shLUC (Sigma-Aldrich, St. Louis, MO, SHC007)

psPAX2 (Addgene, Watertown, MA, Addgene plasmid # 12260)

pMD2.G (Addgene, Watertown, MA, Addgene plasmid # 12259)

Lentiviral plasmid shDCP1a (Origene, Rockville, MD, CAT#: TL305089)

Lentiviral plasmid shDCP1b (Origene, Rockville, MD, CAT#: TL305088)

Lentiviral expression pDCP1b (Origene, Rockville, MD, CAT#: RC207398L1)

### Statistical Analysis

To estimate half-life the equation was used Half-life= -T* ln2/ln(1-NTR), where T is the amount of time that the cells were labeled with 4SU (4 hrs) and NTR is estimated new to total RNA ratio. Z score was used to eliminate outliers (1/3) if needed and at least two half life estimates were used to calculate half life. Variance was calculated and high confidence targets were considered those that had at most 35% variance.

The DDX6 IP had 3 replicas and was performed in control cells (HCT116), DCP1a KO (derived from HCT116), DCP1b KO (derived from HCT116). q-values and p values were calculated. High Confident targets were defined as significant as described above.

DCP1a IP and DCP1b IP each had 2 replicas and were performed in control cells only (HCT116). High Confident targets were defined as significant as described above.

Proteomics LC/MS/MS study of control cells (HCT116), DCP1a KO cells, and DCP1b KO cells was done in triplicates. P-value was calculated. High Confident targets were defined as significant as described above.

The Z score test for two populations, two-tailed, was used to calculate the p value for the comparison of DCP1a/1b dependent mRNAs and mRNAs found in P bodies by Hubstenberger et al.

### Data Availability/Sequence Data Resources

The SLAM-seq raw sequences are available through NCBI SRA (PRJNA1015671). Any dataset not available in the supplementary files can be requested from the corresponding authors.

### Data Availability/Novel Programs, Software, Algorithms

SLAM seq/ GRAND slam programs are available as referenced. Any other script generated during this study can be made available from the corresponding authors on request.

### Web Sites/Data Base Referencing

To define protein domains, we used http://smart.embl-heidelberg.de/. To create Figure 1a we used https://www.biorender.com/.

To perform ontology analysis we used https://www.webgestalt.org/. SLAM-seq QC analysis with MultiQC integrated into SLAM DUNK analysis https://t-neumann.github.io/slamdunk/.

GRAND-SLAM is available at https://github.com/erhard-lab/gedi/wiki/GRAND-SLAM.

## RESULTS

### DCP1a, and not DCP1b, mediates DCP2’s interaction with EDC4 and EDC3

While early eukaryotes express a single DCP1 gene product, the human genome has evolved to encode two paralogs, DCP1a and DCP1b. The advantage of this divergence could relate to the ability to confer tissue-specific roles and expression patterns to the two paralogs. However, expression analyses revealed that DCP1a and DCP1b have concordant rather than reciprocal expression patterns across different mammalian tissues (Supplemental Figure 1). This raises the possibility that DCP1a and DCP1b have evolved to harbor distinct, non-redundant functions in the tissues where they are co-expressed. Understanding any divergent roles for DCP1a and DCP1b requires an understanding of their respective interactions with other components of the mRNA decapping machinery. As mentioned above, yeast have a single DCP1 protein. However, residues that mediate the DCP1 and DCP2 interaction in yeast are not conserved in metazoan cells (31). Yeast DCP2 has a long-disordered C terminal tail that contains 8 low complexity regions which are called short linear motifs (SLiMs)(40) (Figure 1A). DCP2 interacts with other decapping complex members in large part through the avidity effects of SLiMs (40). The C terminal tail of human DCP2 is shorter than that of its yeast counterpart, and some of the SLiMs have been transferred to other subunits of the decapping complex, specifically the enhancers of decapping (EDC) proteins and DCP1a/b (40) (Figure 1A). Both DCP1a and DCP1b also contain low complexity motifs (Figure 1A), suggesting that the human decapping complex has undergone significant re-wiring, when compared to yeast (40). As such, mechanisms mediating assembly and integrity of the yeast complex provide little information relevant to the human complex. As a key example, it is unknown which of the interactions between the primary members of the decapping complex, if any, are supported by DCP1a versus DCP1b. To directly assess the relative roles of DCP1a and DCP1b in decapping complex assembly, we immunoprecipitated DCP2 from the human colorectal carcinoma line HCT116. Results from parental cells were compared with results from HCT116 cells that had been depleted of either DCP1a or DCP1b. These studies demonstrated that depletion of DCP1a, but not DCP1b, resulted in a decrease in the interaction between DCP2 and EDC4 (Figure 1B). While this finding is consistent with DCP1a and DCP1b having distinct roles in the decapping complex, it also remained possible that DCP1a and DCP1b have similar functions, but that cellular DCP1B expression levels were too low to compensate for the depletion of DCP1a. To assess this, DCP1b was ectopically expressed at elevated levels in DCP1a-depleted cells. Immunoprecipitation of DCP2 from DCP1a-depleted cells showed that ectopic overexpression of DCP1b failed to rescue the interaction between DCP2 and EDC4 (Figure 1B), providing additional support for a model in which DCP1a and DCP1b have intrinsically distinct functions that are unrelated to their relative expression levels.

In the shRNA studies described above, endogenous DCP1a and DCP1b levels were reduced by approximately 80%. While informative, it was also important to assess the differential roles of DCP1a and DCP1b in cells where the genes encoding these were deleted. For this purpose, single cell clones lacking DCP1a (DCP1a KO) or DCP1b (DCP1b KO) were isolated, and loss of protein confirmed (Figure 1C). As in cells where depletion was induced using shRNA, complete deletion of DCP1a also resulted in a defect in the interaction between DCP2 and EDC4, while DCP1b deletion had no effect on this interaction (Figure 1C). Consistent with this, the decapping complex subunit EDC3 was similarly impacted by DCP1a loss, but not by DCP1b loss (Figure 1C). As in shRNA-treated cells, ectopic DCP1b overexpression did not rescue these interactions (Figure 1D), again supporting a model in which DCP1a and DCP1b have qualitatively distinct roles in the assembly/integrity of the mRNA decapping complex.

### DCP1a is critical for interactions between known decapping complex members

Since targeted analysis revealed that loss of DCP1a and DCP1b have distinct effects on the interaction of DCP2 with EDC3 and EDC4, it was important to more broadly examine the role of DCP1a/b in the decapping complex interactome. To address these effects, the interactome of the RNA helicase DDX6 was examined using a proteomic approach. (DDX6 interacts with DCP1a, DCP1b, and DCP2, thus allowing interrogation of decapping complex integrity without disturbing interactions among the core complex members.)

#### Immunoprecipitation and quantitative

LC/MS/MS analysis identified 2069 DDX6 interactors. The interactomes of DDX6 varied depending on the presence or absence of DCP1a or DCP1b (Figure 2A). In parental cells, DDX6 interacts with EDC3, EDC4, DCP1a, and DCP1b, with no DCP2 being detected (Figure 2B). Deletion of DCP1a reduced the interaction of both EDC3 and EDC4 with DDX6 (Figure 2B). Conversely, the DDX6-EDC4 interaction was enhanced in DCP1b-depleted cells (Figure 2B). These findings are consistent with the empirical immunoprecipitation data shown in Figure 1. One interpretation of these findings is that DCP1a acts as a positive regulator of decapping complex assembly. The role of DCP1b in assembly of the decapping complex is multifaceted, with no discernable impact on the DDX6-EDC3 interaction, but a negative impact on DDX6-EDC4 interactions.

A core set of 53 proteins interact with DDX6 regardless of DCP1a or DCP1b loss (Figure 2A). In broad terms, DCP1a loss decreased the interactions of DDX6 with other RNA regulating proteins whereas DCP1b loss did not (Figure 2C). Ontology analysis showed the core 53 proteins are enriched in PB & stress granule proteins, telomere regulating proteins, and proteins with demethylase activity (Figure 2D). We identified 6 proteins whose interaction with DDX6 depends on DCP1a, 20 proteins whose interaction depends on DCP1b and 18 proteins whose interaction with DDX6 depends on both DCP1a and DCP1b (Figure 2A). Ontology analysis revealed that DCP1a-dependent proteins are enriched for proteins that regulate synaptic vesicle exocytosis, deadenylation-dependent and -independent decapping of mRNA, and mRNA cap binding (Figure 2E). One of the most enriched categories within DCP1b dependent proteins are those involved in translation initiation (Figure 2E). Both DCP1a-and DCP1b-dependent proteins participate in regulating mRNA splicing and telomerase RNA binding (Figure 2E).

### DCP1a and DCP1b have primarily non-overlapping interactomes

To further elucidate the multi-faceted roles of DCP1a and DCP1b in the decapping complex and elsewhere, we immunoprecipitated DCP1a and DCP1b from parental HCT116 cell lines. LC-MS/MS analysis of these reactions revealed that DCP1a interacts at high confidence with 103 proteins and DCP1b with 705 proteins, where 11 proteins are found in both IPs and include core decapping complex members DCP1a, DCP1b, and EDC3. Not all 103 and 705 proteins in the DCP1a/b immunoprecipitations are direct binding partners. Ontology analysis revealed that DCP1a interacts with proteins that have histone and protein deacetylase activity, bind mRNA, tubulin, and chromatin, and have transcription corepressor activity (Figure 3A). Cellular component analysis shows that DCP1a can be found in P bodies but it also in histone deacetylase complexes, and spindle, chromatin and oxoglutarate dehydrogenase complexes (Figure 3A). In contrast, the DCP1b interactome is enriched in proteins that regulate translation, including many proteins that have aminoacyl-tRNA ligase activity (Figure 3B). However, DCP1b also interacts with proteins that bind cadherin, RNA, and purine ribonucleotides (Figure 3B). DCP1b also interacts with proteins involved in neutrophil activation, translational initiation, and focal adhesion/adherens junctions (Figure 3B). These data and previous studies suggest that while DCP1a and DCP1b interact with and bind RNA, they have other cellular functions. Based on this ontology analysis of DCP1a and DCP1b interactomes (Figure 3), and the ontology analysis of the DCP1a and DCP1b dependent proteins in DDX6 IP (Figure 2), DCP1a appears to play a disproportionate role in transcription and DCP1b in translation.

**Figure 3.**
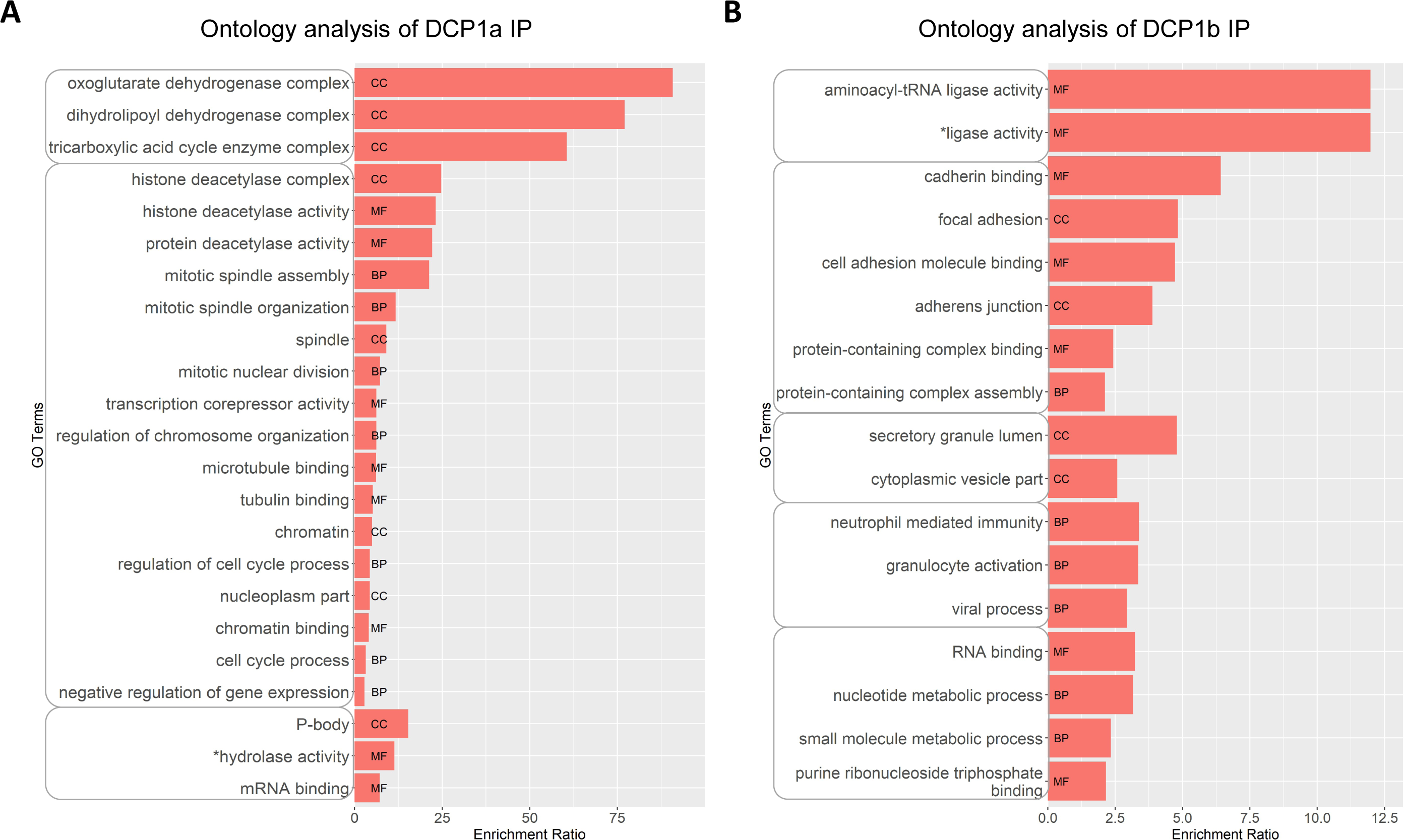
DCP1a and DCP1b have primarily non-overlapping interactomes. **A)** DCP1a was immunoprecipitated from HCT116 cell lysates and the interactome profiled by LC/MS/MS. Ontology analysis of the proteins associated with DCP1a is shown. Only relevant GO terms with p value < 0.05 are shown. GO terms with similar cellular role are grouped. MF-Molecular function, BP – Biological process, CC – Cellular Compartment. *GO terms have been edited for brevity in some cases. Full GO terms can be found in supplemental data. **B)** DCP1b was immunoprecipitated from parental HCT116 cells and the interactome profiled by LC/MS/MS. Ontology analysis of the proteins associated with DCP1b is shown. Only relevant GO terms with p value < 0.05 are shown. GO terms with similar cellular role are enclosed. MF-Molecular function, BP – Biological process, CC – Cellular Compartment. *GO terms have been edited for brevity in some cases. Full GO terms can be found in supplemental data.

### The impact of DCP1a and DCP1b on mRNA turnover

EDC3 and EDC4 regulate the efficiency and kinetics of decapping (41,42), and DCP2 can impact the degradation rates of specific groups of mRNAs (43–45). Since DCP1a and DCP1b impact decapping complex interactions involving EDC3, EDC4, and other subunits, experiments were designed to examine their impact on the half-lives of specific groups of mRNAs. To examine the effects that DCP1a and DCP1b have on global RNA half-life, *thiol(SH)-linked alkylation for the metabolic sequencing of RNA* (SLAM-seq) was performed (37). In SLAM-seq, transcriptome-wide RNA sequencing detects 4 thiouridine (4SU) incorporation into RNA at single-nucleotide resolution, based on T > C conversions (37) (Figure 4A). Conditions were optimized for this experiment by evaluating the impact of different 4SU concentrations on cell viability (Supplemental Figure 2). Parental, DCP1a KO and DCP1b KO cells were cultured with 4SU and cellular uptake of 4SU was confirmed by dot blot analysis of 4SU-incorporation into RNA (Supplemental figure 3). SLAM-DUNK (37) and GRAND-SLAM algorithms (36) were utilized to analyze the mRNA libraries, and the MultiQC module of SLAM-DUNK was used to assess the TC conversion rates, normalized across UTRs (Figure 4B). Half-life was calculated for 4,439 mRNA molecules in control cells, 4,018 in DCP1a KO cells, and 3,328 in DCP1b KO cells. The impact of DCP1a and DCP1b on mRNA half-life regulation was examined by analyzing 1,440 mRNAs for which a half-life was calculated in all three conditions (Figure 4C). Of these, 734 mRNAs were not significantly impacted by the loss of either DCP1a or DCP1b. Based on the assumption that loss of DCP1a/b should decrease mRNA decapping and thereby increase mRNA stability, direct mRNA targets are predicted to have significantly longer half-lives in the absence of DCP1a or DCP1b. Using this criteria, 131 mRNAs are controlled by DCP1a alone, 309 by DCP1b alone, and 140 mRNAs were controlled by both DCP1a and DCP1b.

**Figure 4.**
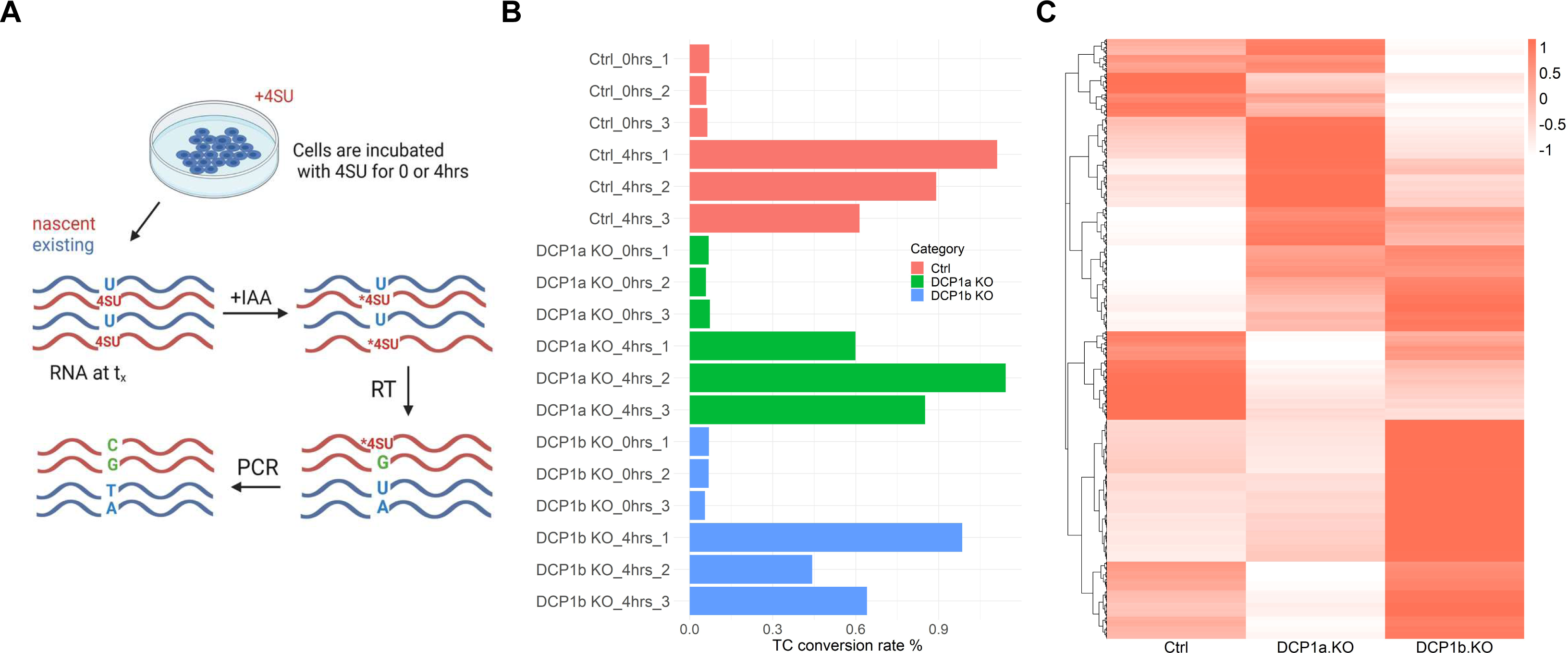
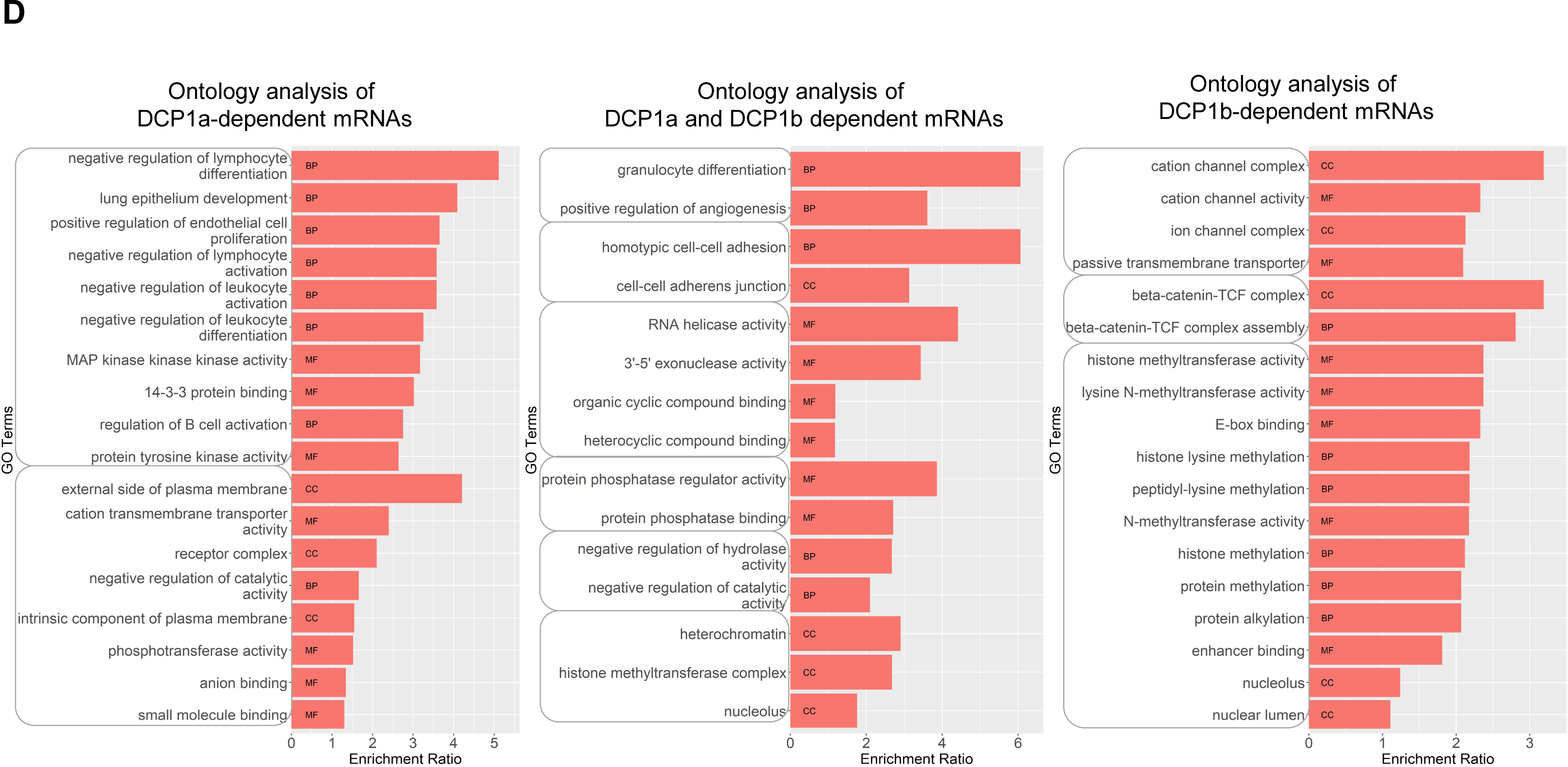
DCP1a and DCP1b have differential impacts on mRNA turnover. **A)** Experimental pipeline for determining mRNA t_1/2_ using thiol (SH)-linked alkylation for the metabolic sequencing of RNA (SLAM-seq). This method allows direct quantification of 4-thiouridine (4SU) within the 4SU labeled mRNA. Control, DCP1a KO and DCP1b KO cells were cultured with 100 µM 4SU for 0 or 4 hours. Cells were harvested, RNA was isolated and alkylated, under conditions of light restriction. Alkylated RNA was purified, mRNA libraries were prepared and subjected to RNA-seq. When 4SU, as an uracil analog, is incorporated and alkylated, reverse transcriptase misincorporates guanosine. The 4SU content of mRNA was quantified by measuring T > C conversion in the final 3’ end mRNA sequencing. **B)** Overall T>C conversion efficiency at 0hr and 4hr of 4SU incubation in ctrl, DCP1a KO, and DCP1b KO cells. Three biological replicates (1,2 and 3) are shown. **C)** Heatmap representing average t_1/2_ of 1440 mRNAs in ctrl, DCP1a KO and DCP1b KO cells. The heatmap is scaled and clustered by rows. **D)** Upon loss of DCP1a, DCP1b or both, t_1/2_ of certain groups of mRNAs are upregulated. These mRNAs were designated DCP1a-, DCP1b- or DCP1a and DCP1b dependent mRNAs. Ontology analysis of 271 mRNAs dependent on DCP1a, 140 mRNAs dependent on DCP1a and DCP1b, and 449 mRNAs dependent on DCP1b, is shown. GO terms with similar cellular role are grouped. Only relevant GO terms with p value < 0.05 are shown. MF-Molecular function, BP – Biological process, CC – Cellular Compartment.

Ontology analysis of the mRNAs controlled by DCP1a and/or DCP1b was carried out using WebGestalt (38) with results demonstrating that all three categories of transcripts are enriched for those encoding histone methyltransferase complex subunits, cell adhesion molecules and proteins linked to RNA helicase activity (Figure 4D). The 271 mRNAs specifically regulated by DCP1a are enriched for those encoding proteins involved in transmembrane transport, the beta-catenin TCF complex, and cell differentiation, while the 449 mRNAs controlled by DCP1b are enriched for those encoding proteins regulating lymphocyte, leucocyte, fat cell differentiation, and transmembrane transport activity (Figure 4D). These findings add further support to the concept that DCP1a and DCP1b are functionally distinct.

### The impact of DCP1a and DCP1b on mRNA half-life is buffered through their effect on transcription

The decapping complex has been implicated in transcript buffering, a process in which mRNA levels are kept constant through reciprocal changes in mRNA synthesis and degradation (46–48). Since DCP1a and DCP1b altered the half-lives of specific RNAs, protein levels encoded by those RNA were assessed. For this purpose, protein lysates were generated from parental, DCP1a KO, and DCP1b KO cells and then subjected to quantitative profiling by LC/MS/MS. A total of 5,543 proteins were identified. Proteins identified in DCP1a KO and DCP1b KO were compared to those identified in parental cells. 869 proteins were identified as high confidence in DCP1b KO, 741 proteins in DCP1a KO, and 238 high confidence in both DCP1a KO and DCP1b KO cells. Most of these proteins in DCP1a KO and DCP1b KO cells had a log2 ratio of less than 1 or higher than -1, effectively they were unaltered (Figures 5A-B). 44 proteins in DCP1a KO (Figure 5A) and 38 proteins in DCP1b KO have a log2 ratio greater than 1 or lower than -1, when compared to parental cells (Figure 5B). DCP1a loss significantly impacted proteins that bind mRNAs and have telomerase inhibitor activity (Figure 5C). Conversely, DCP1a loss significantly impacted levels of proteins that bind dynein/myosin/actin and those playing a role in cell-cell adhesion mediated by integrin (Figure 5D).

**Figure 5.**
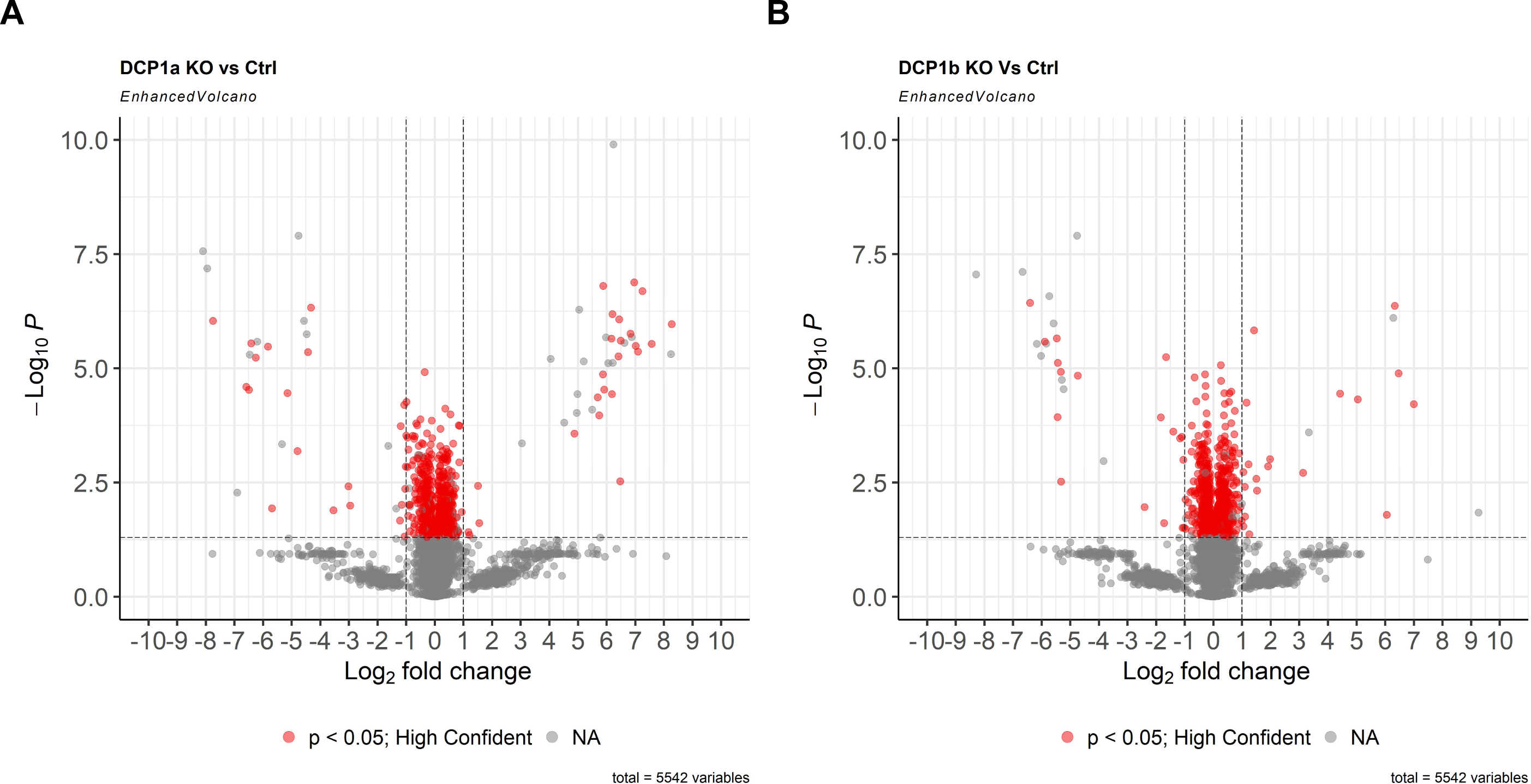
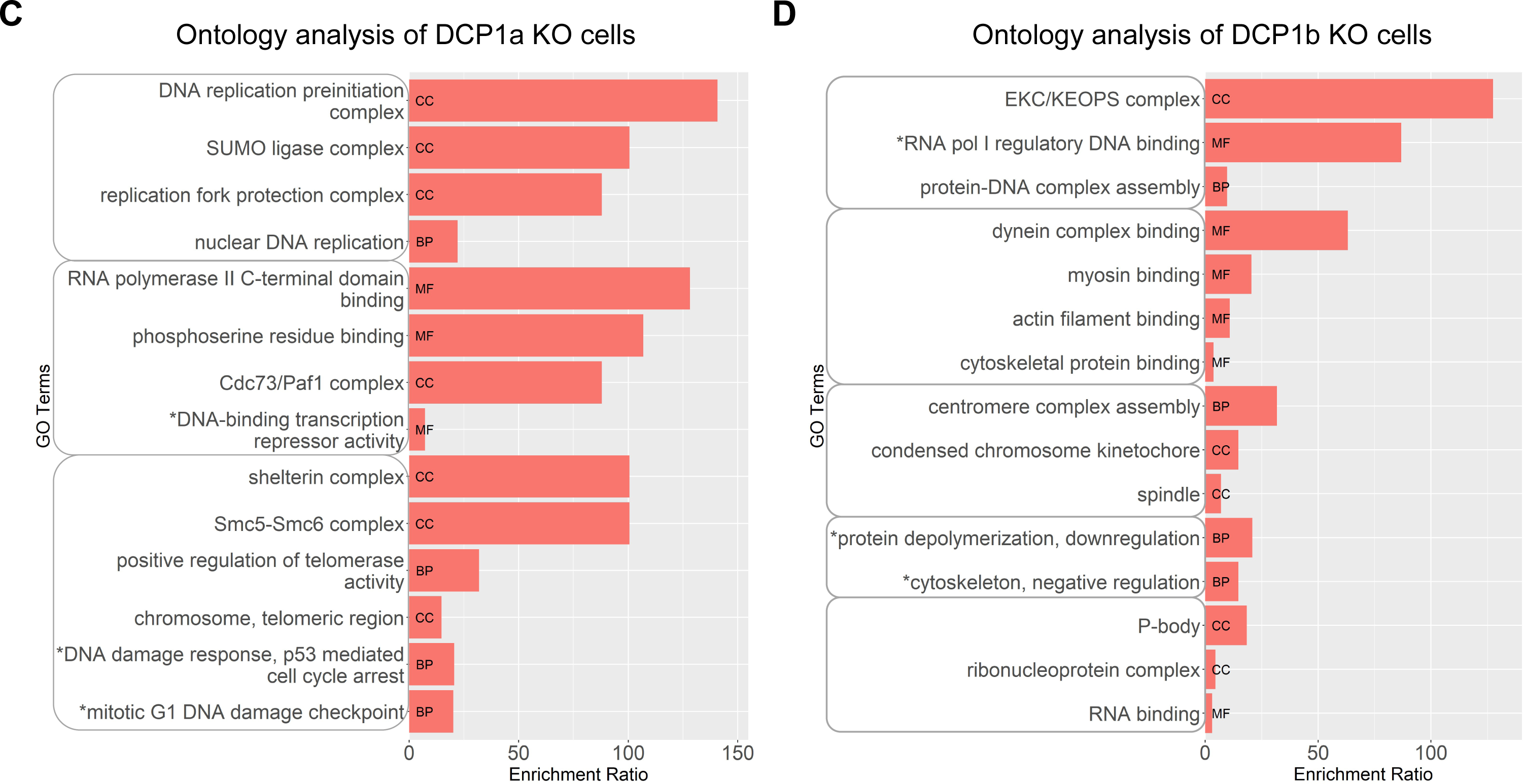
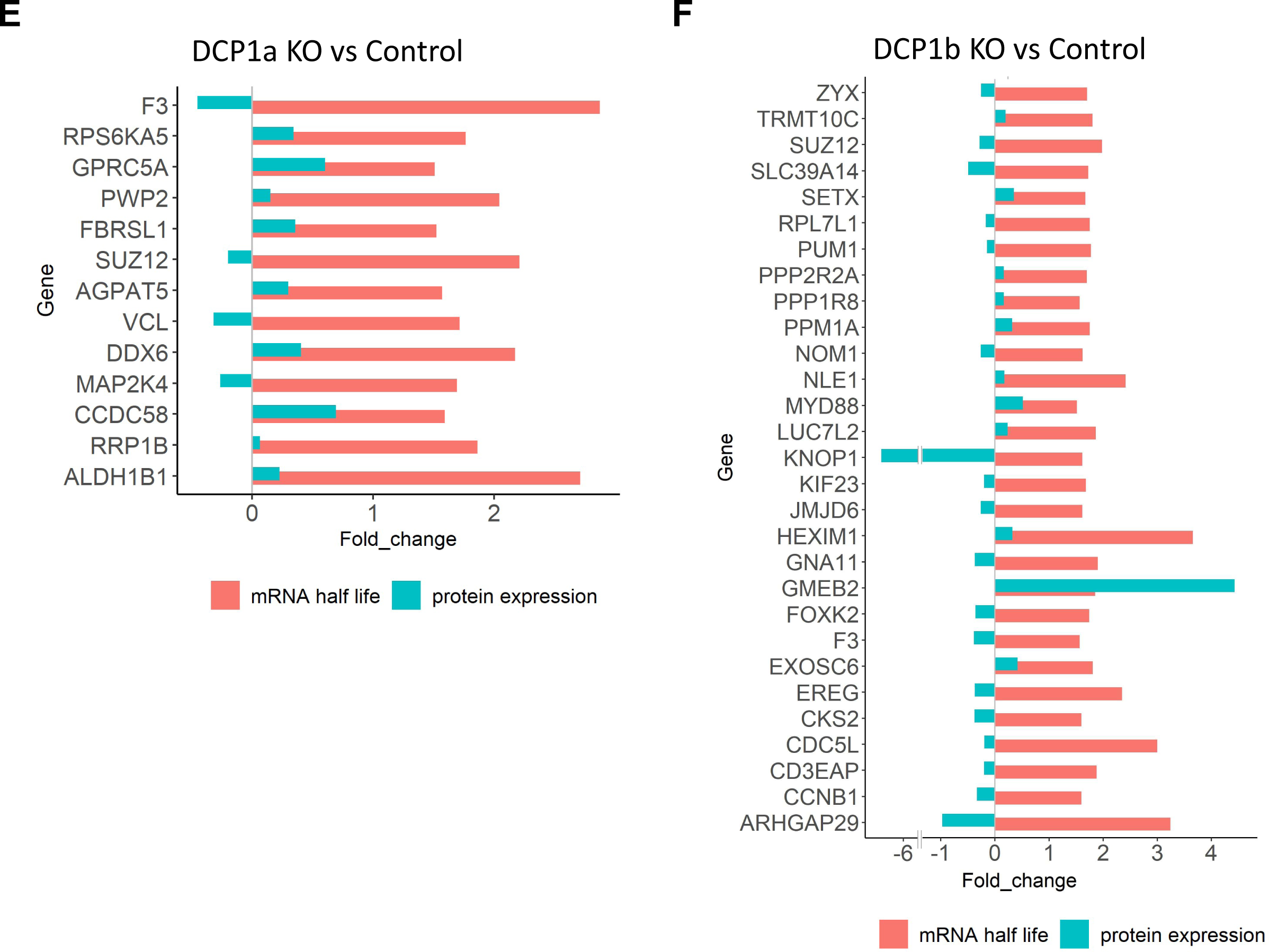
The impact of DCP1a and DCP1b on mRNA t_1/2_ is buffered via effects on transcription. **A)** Lysates were collected from control, DCP1a KO and DCP1b KO cells and subjected to SDS/PAGE. Proteins were stained, bands excised and analyzed by LC/MS/MS. Volcano plot shows log2(Fold change) vs log10(p value) in DCP1a vs ctrl condition protein levels of the 5542 proteins identified. The vertical line indicates log2-fold change of 1 or -1. The horizontal line demarcates a p value of 0.05. Data points with p<0.05 were deemed to be of high confidence and are shown in purple. **B)** Lysates were collected from control, DCP1a KO and DCP1b KO cells and analyzed as in (A). Volcano plot shows log2(Fold change) vs log10(p value) in DCP1b vs ctrl condition protein levels of the 5542 proteins identified. **C)** Ontology analysis of only significantly impacted proteins (which did not appear in DCP1b KO results) in DCP1a KO vs ctrl. GO terms with similar cellular role are grouped. Only relevant GO terms with p value < 0.05 are shown. MF-Molecular function, BP – Biological process, CC – Cellular Compartment. **D)** Ontology analysis of only significantly impacted proteins (which did not appear in DCP1a KO results) in DCP1b KO vs ctrl. **E)** Comparison of log2 fold change in protein level and t_1/2_ of DCP1a-dependent mRNAs. **F)** Comparison of t_1/2_ of DCP1b-dependent mRNAs and log2 fold change in protein level.

The proteomic and SLAM-seq datasets were compared to specifically assess whether transcripts with increased half-life had corresponding increases in protein levels. This analysis revealed that protein levels for those mRNAs whose half-lives are upregulated, are not significantly impacted. Specifically, from a total 271 targets whose half-lives were upregulated by DCP1a loss, 109 are identified in the proteomics dataset, but only 13 are statistically significant in the proteomics dataset. However, these 13 proteins were not upregulated on protein level (Figure 5E). Of 449 targets whose half-lives are upregulated upon DCP1b loss, 190 are identified but only 29 are statistically significant in the proteomics dataset. Among these 29 proteins, one was upregulated, one was downregulated, and 27 were not impacted on protein level (Figure 5F). The lack of congruence between mRNA half-life and protein levels reflects translational repression of these specific mRNAs.

## Discussion

Regulation of mRNA decapping plays a critical role in gene expression. A single DCP1 co-factor modulates activity of the decapping complex catalytic subunit DCP2 in lower eukaryotes such as yeast. In higher eukaryotes, including humans, two paralogous cofactor proteins, DCP1a and DCP1b, work with DCP2. The studies reported here represent the first to define the key functional distinctions between the human DCP1 paralogs. Despite both containing the conserved protein domain (EVH1), DCP1a and DCP1b differ in terms of protein interactomes and in their role controlling integrity of the mRNA decapping complex itself (Figures 1, 2, 3). DCP1a regulates RNA cap binding proteins association with the decapping complex and DCP1b controls translational initiation factors interactions (Figure 2E). Functionally, the transcripts impacted by DCP1a loss are distinct from those impacted by DCP1b loss (Figure 4). Specifically, DCP1a is predominantly involved in regulation of transcription pathways while DCP1b engages in regulation of translation pathways. Finally, our data underscore the role of the decapping complex in transcript buffering (Figure 5). Collectively, these findings demonstrate that DCP1a and DCP1b function as distinct, non-redundant cofactors of the decapping enzyme DCP2.

As mentioned above, DCP1a and DCP1b have unique roles in decapping complex integrity. For example, DCP1a impacts the interaction of DCP2 with EDC4 and EDC3 (Figure 1B, 1C, 1D and 2B), in a manner that cannot be compensated for by ectopic DCP1b expression (Figure 1B and 1D). On the other hand, DCP1b plays an inhibitory role in the recruitment of EDC4 (Figure 2B). This same pattern is observed when evaluating decapping complex interactions more broadly, where DCP1a depletion reduces the intensity of the DDX6 interactions while DCP1b depletion enhances it (Figure 2C). One interpretation of these data is that DCP1a and DCP1b have reciprocal roles in decapping complex assembly/integrity, with DCP1a functioning as a positive regulator and DCP1b as a negative regulator. Since DCP1a and DCP1b can form either homo- or heterotrimers within the decapping complex, modulating the stoichiometry of DCP1a-DCP1b content could provide a cellular mechanism for regulating complex assembly and decapping activity.

As shown, DCP1a and DCP1b impact the decay rates of distinct groups of mRNAs. Specifically, we defined 271 DCP1a-dependent mRNAs and 449 DCP1b dependent mRNAs (Figure 4). Initial studies identified no primary sequence motifs that correlated with DCP1a or DCP1b dependence. Overrepresented among the transcripts controlled by DCP1b, and to a lesser extent DCP1a, are those encoding subunits of histone methyltransferase complexes involved in transcriptional regulation (Figure 4D). The pool of DCP1a-dependent mRNAs encode proteins regulating lymphocyte differentiation and activation. As these studies were conducted in the colorectal carcinoma line HCT116, the physiological implications of this impact on lymphocyte-associated transcripts is unclear. While this study represents the first to define DCP1a- and DCP1b-dependent mRNAs, the ontology analysis is consistent with previous reports implicating DCP1a in the immune response and the decapping complex in transcription regulation (49,50).

It is important to note that our data may be biased by preferentially including highly transcribed mRNAs. The SLAM-Seq methodology relies on the conservative use of 4SU, which dramatically impacts cell stress signaling when used at higher concentrations. The low concentrations of 4SU used here allowed the analysis of 1440 DCP1a or DCP1b-regulated transcripts, presumably because those are transcribed at a rate high enough to incorporate significant levels of 4SU during the labeling interval.

Somewhat paradoxically, the levels of specific mRNA molecules regulated by DCP1a and DCP1b did not universally correlate with the level of the proteins encoded by these transcripts (Figure 5). mRNA decapping complex members have previously been implicated in establishing a transcript feedback loop that “buffers” steady state protein levels, even in the face of alterations in mRNA. Related to this phenomenon, some transcripts localized to P bodies are maintained in a state of translational repression. This prompted a comparison between the DCP1a- and DCP1b-dependent transcripts defined here, and those known to be enriched in P bodies (21). This analysis revealed that most DCP1a- and DCP1b-dependent transcripts are among those previously reported to localize to P bodies, 165/271 (61%) and 273/449 (61%) respectively (p value < 0.001). Similarly, 88/140 (63%) of the targets impacted by both DCP1a and DCP1b are localized to P bodies (p value < 0.001). Considered together, these findings support the developing model that mRNA decapping is a key element in the transcript buffering pathway. Also supporting this concept is our discovery that DCP1a interacts closely with proteins involved in transcription (Figure 3A), and DCP1b interacts closely with the translation machinery (Figure 3B), while also regulating stability of mRNAs involved in transcription (Figure 4D) and being localized to nucleus and cytoplasm (Supplemental Figure 4).

The ability of DCP1a and DCP1b to differentially influence the overall composition of the mRNA decapping complex provides an opportunity to differentially control distinct sets of mRNA targets. It also provides a mechanism for controlling the distinct functions of the decapping complex, e.g., mRNA stability, translation efficiency, and/or transcript buffering. From the initial studies reported here, it is clear that DCP1a is uniquely essential for efficient decapping complex assembly, interacting with RNA cap binding proteins, as well as being responsible for the turnover of mRNAs which are directly involved in both adaptive immunity and transcription. In contrast, DCP1b plays a unique role in fostering the interaction of the decapping complex with the translational machinery, while simultaneously playing a role in the turnover of mRNAs involved in transcription.

Ultimately it will be important to delineate how the relative stoichiometry of DCP1a and DCP1b in the decapping complex is controlled when cells are exposed to different stimuli, and how altering that stoichiometry impacts decapping activity and specificity. As the biochemical events responsible for transcript buffering are elucidated, it will also be critical to understand how DCP1a and DCP1b modulate those events.

### Data Availability/Sequence Data Resources

The SLAM-seq raw sequences are available through NCBI SRA (PRJNA1015671). Any dataset, not included in the supplemetary files, is available from the corresponding authors on request.

## Acknowledgements

We thank Andrew Kossenkov Ph.D. (Bioinformatics Facility, Wistar Institute), Hsin-Yao Tang Ph.D. (Proteomics Facility, Wistar Institute), Sonali Majumdar M.S., (Genomics Facility, Wistar Institute), Fadia Ibrahim Ph.D.(Thomas Jefferson University), Tobias Neumann, Ph.D., Florian Erhard Ph.D., Alison Moss, Ph.D., Yohei Kirino, PhD, Mike Kiledjian Ph.D., Alexander Mazo, Ph.D., Emad Alnemri, Ph.D., and Amanda Oran Ph.D.

---

**Supplemental Figure 1.**
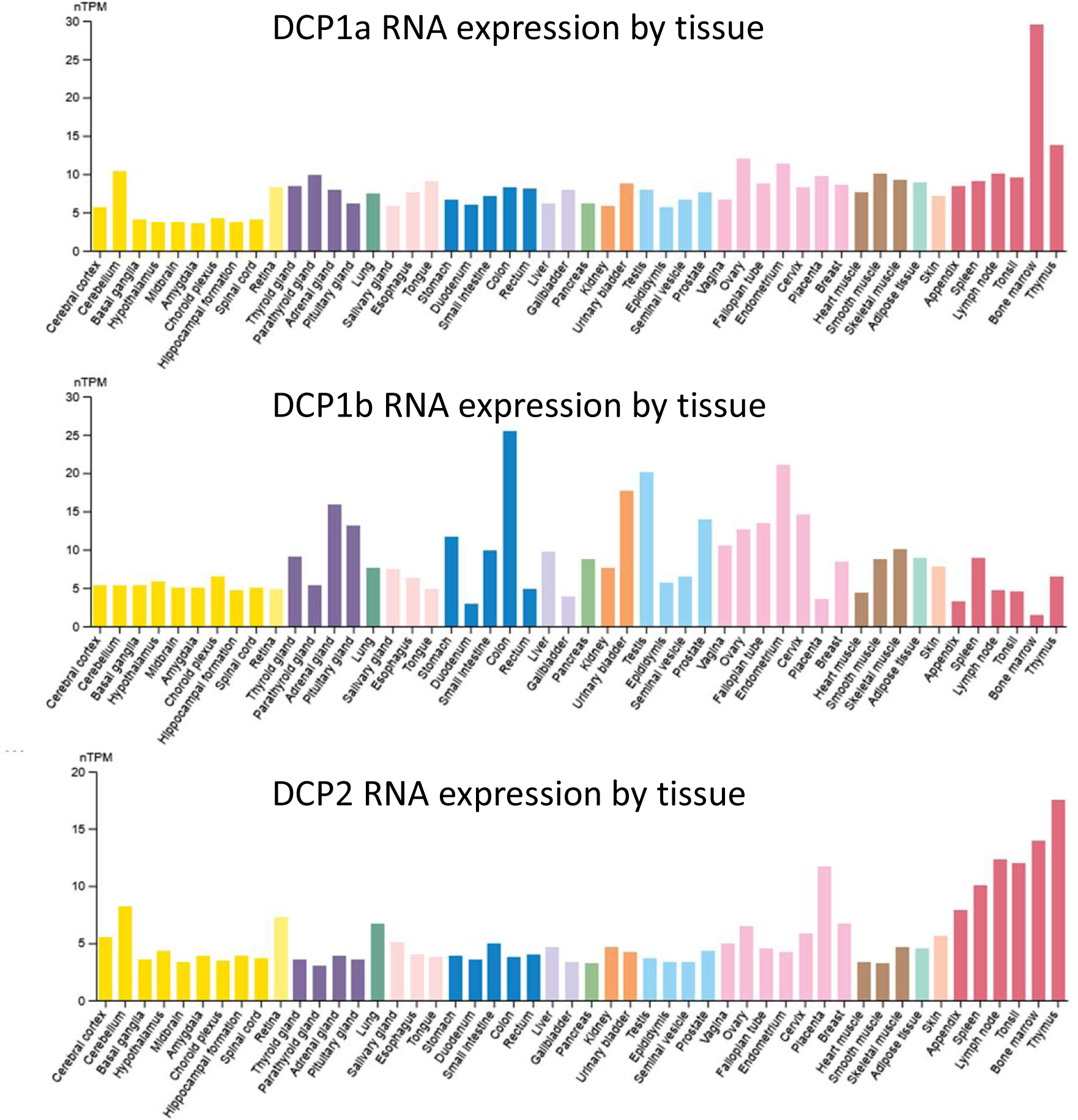
DCP1a, DCP1b and DCP2 RNA expression by tissue. The graphs are from www.proteinatlas.org. This is the consensus dataset consisting of normalized RNA expression (nTPM) levels for 55 tissue types.

**Supplemental Figure 2.**
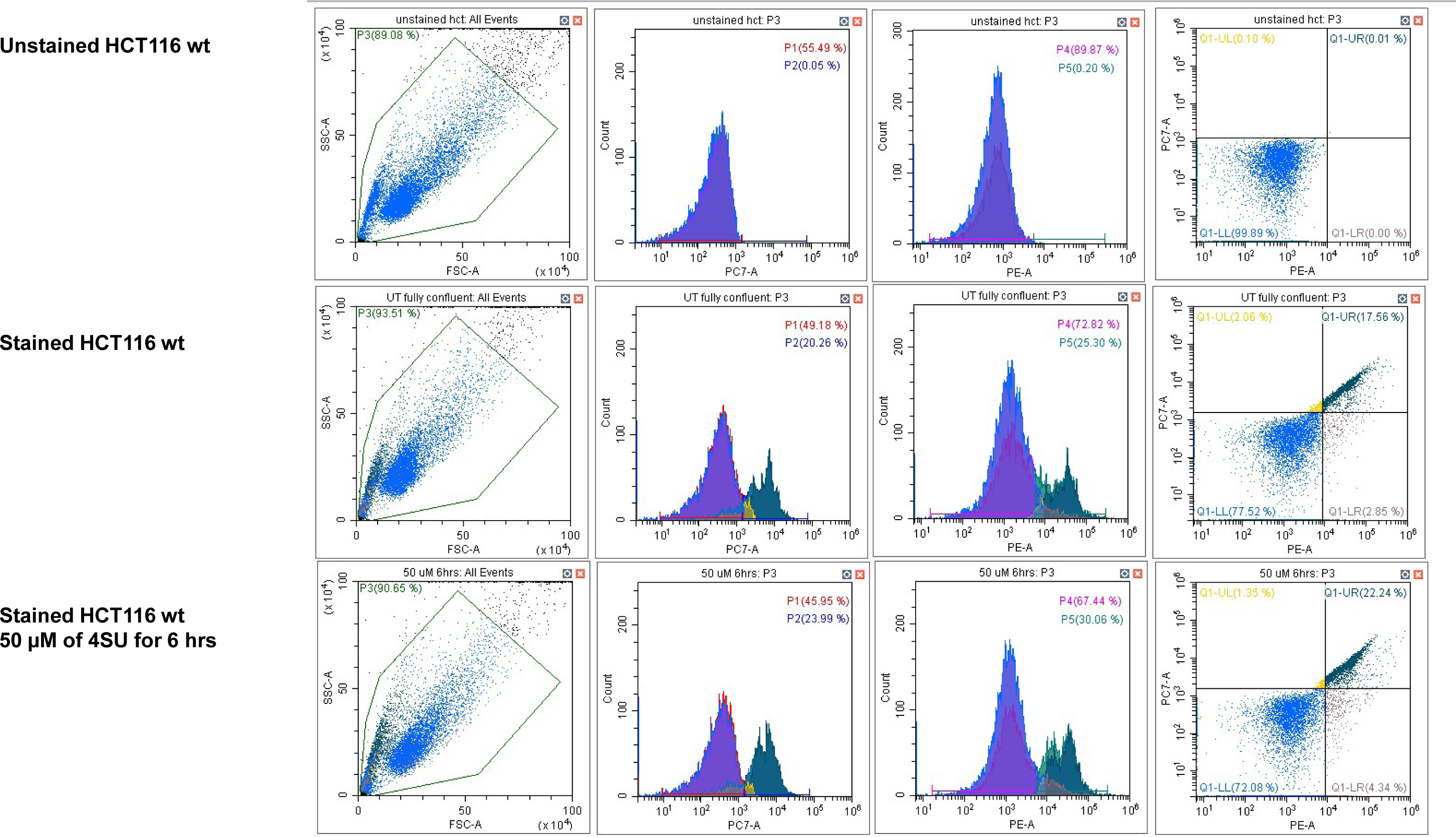
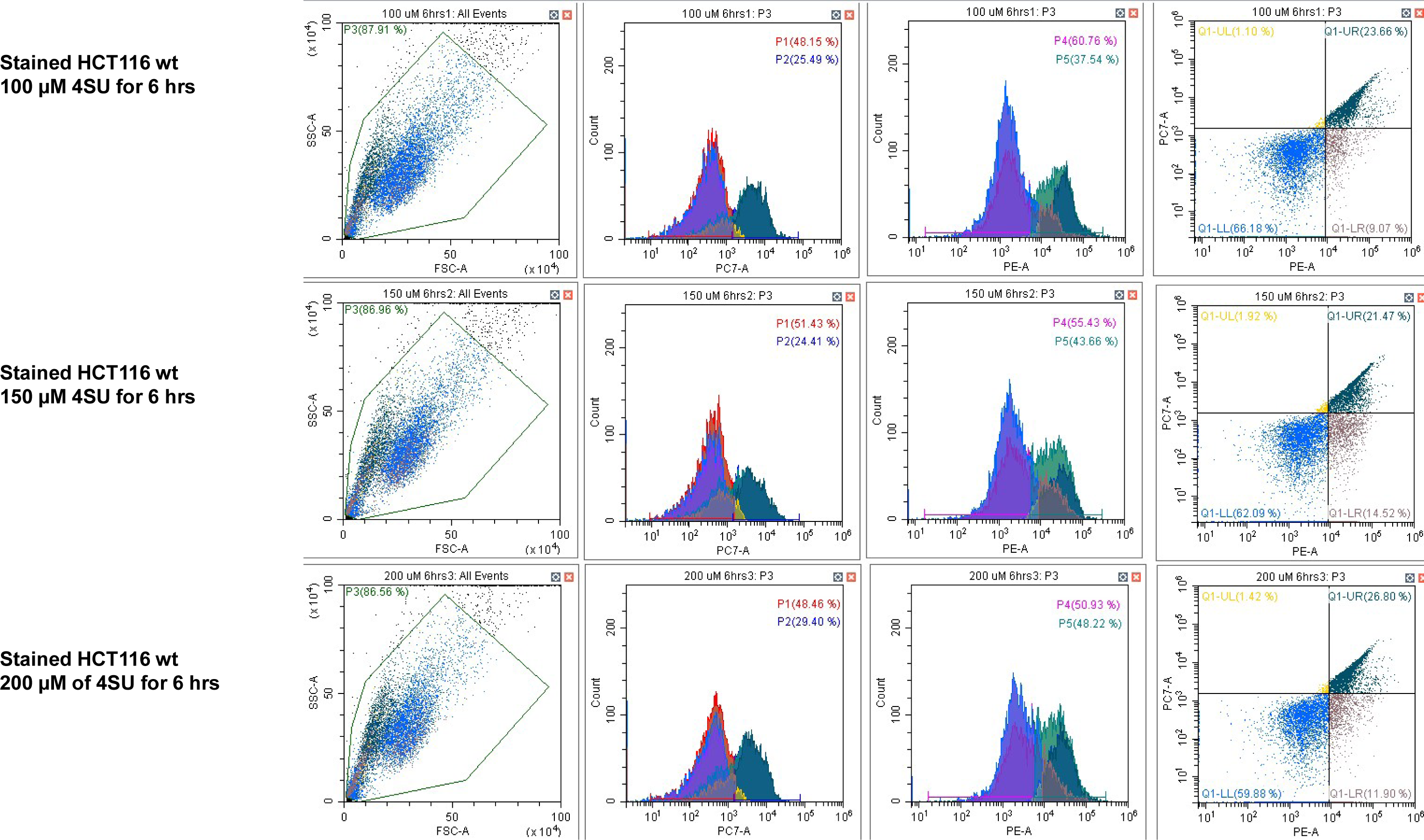
Evaluation of 4SU cell toxicity. Cells were cultured to an appropriate confluence and media containing 0 µM, 50 µM, 100 µM or 200 µM 4SU was added. Cells were cultured in the presence of 4SU for 6hrs while supplying fresh 4SU media every 3 hrs. At 6hrs, media was replaced with DMEM without 4SU and 24hrs later cell viability was evaluated. Cells were stained with either nothing or with propidium iodide (PI) and/or Annexin V. Cell death was assessed with *CytoFLEX S Flow cytometer*.

**Supplemental Figure 3.**
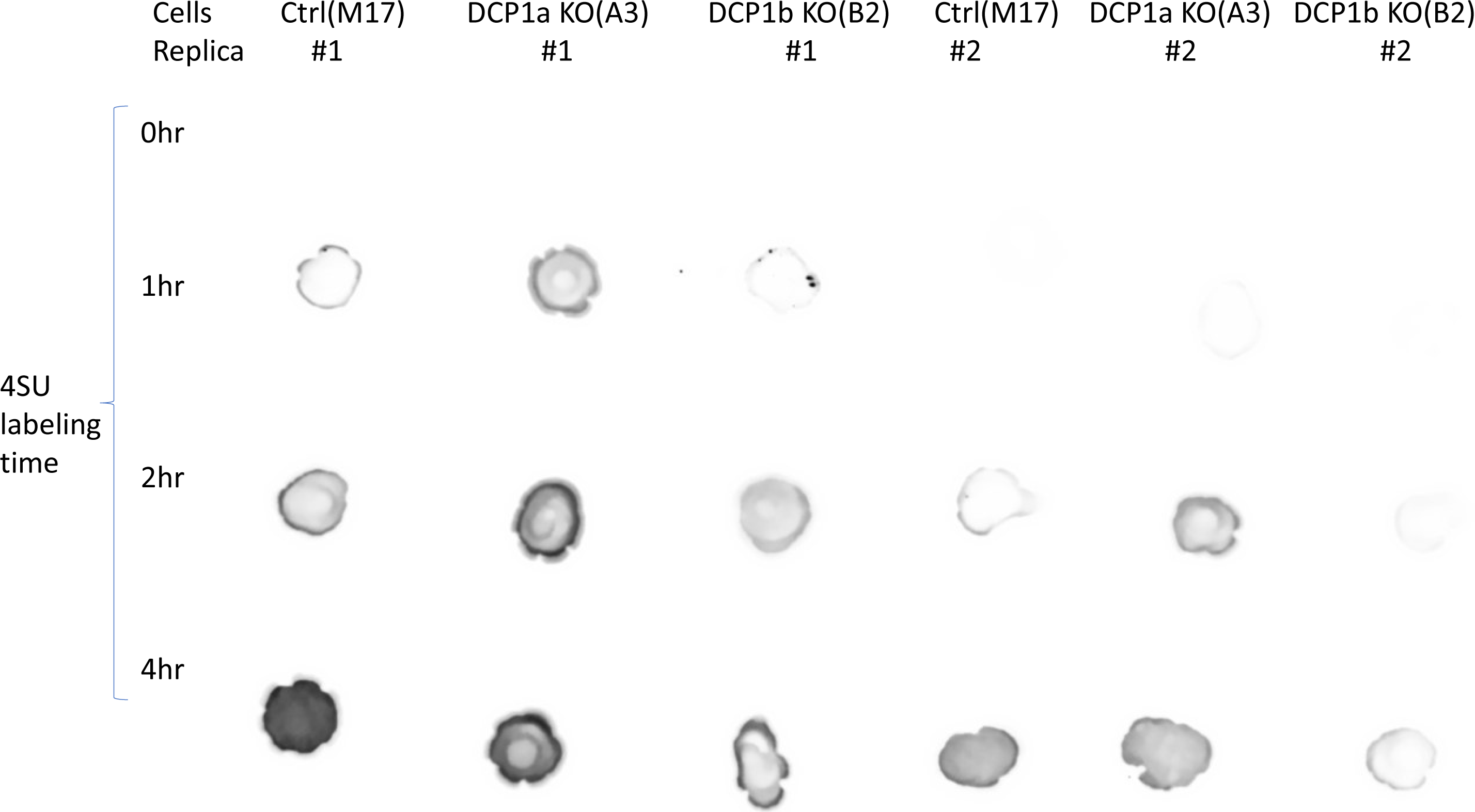
Evaluation of 4SU cell uptake. 4SU incorporation was confirmed with dot blot analysis of 4SU labeled RNA, prior to alkylation, in control cells, DCP1a KO and DCP1b KO cells. Replicas 1, and 2 are shown for 0,1,2 and 4hr incubation with 4SU media.

**Supplemental Figure 4.**
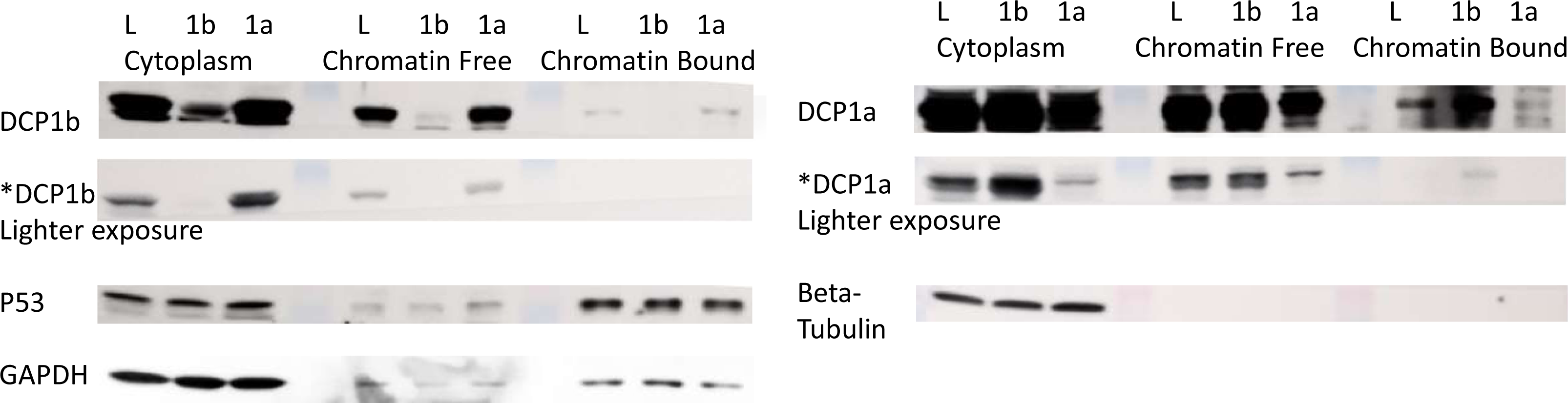
DCP1a and DCP1b are found in chromatin bound and chromatin free cellular extracts. HCT116 cells were infected with viruses targeting either shLUC (L), shDCP1a (1a), or shDCP1b (1b). Cellular fractionation was performed.

